# Click evoked middle ear muscle reflex: Spectral and temporal aspects^a)^

**DOI:** 10.1101/2020.08.24.265462

**Authors:** Sriram Boothalingam, Shawn S. Goodman

**Author notes:** Portions of this work were presented at the 47th Annual Scientific & Technology Conference of the American Auditory Society, Scottsdale, AZ, March 2020.

## Abstract

This study describes a time series-based method of middle ear muscle reflex (MEMR) detection using bilateral clicks. Although many methods can detect changes in the OAE evoking stimulus to monitor the MEMR, they do not discriminate between true MEMR-mediated vs. artifactual changes in the stimulus. We measured MEMR in 20 young clinically normal hearing individuals using 1-second-long click trains presented at six levels (65 to 95 dB peak-to-peak SPL in 6 dB steps). Changes in the stimulus levels over the 1 second period were well-approximated by two-term exponential functions. The magnitude of ear canal pressure changes due to MEMR increased monotonically as a function of click level but non-monotonically with frequency when separated into 1/3^rd^-octave wide bands between 1 and 3.2 kHz. MEMR thresholds estimated using this method were lower than that obtained from a clinical tympanometer in ∼94% of the participants. A time series-based method, along with statistical tests, may provide additional confidence in detecting the MEMR. MEMR effects were smallest at 2 kHz, between 1 and 3.2 kHz, which may provide avenues for minimizing the MEMR influence while measuring other responses (e.g., the medial olivocochlear reflex).

## I. INTRODUCTION

Two auditory reflexes exert influence on peripheral auditory signal processing. The middle ear muscle reflex (MEMR) influences signal transfer by altering the impedance characteristics of the middle ear, and the medial olivocochlear reflex (MOCR) inhibits the cochlear active process (review: Guinan, 2006). Animal models suggest both of these processes benefit the organism by protecting cochlear structures from loud sounds and improving signal-to-noise ratio (Liberman & Guinan, 1998; see also Kirk & Smith, 2003). As such, understanding the functioning of these two reflexes in humans holds both scientific and clinical value.

Otoacoustic emissions (OAE)-based assays provide a convenient and non-invasive means to investigate MOCR effects in humans. However, many of the stimuli which activate the MOCR also simultaneously activate the MEMR. Both the input to cochlea (stimulus) and the retrograde emissions can be significantly altered by MEMR activation (Guinan, Backus, Lilaonitkul, & Aharonson, 2003). Thus, a major impediment to measuring MOCR is that, when active, the MEMR effects on OAEs can masquerade as MOCR effects (Goodman, Mertes, Lewis, & Weissbeck, 2013; Goodman, Venkitakrishnan, Adkins, & Mueldener, 2018; Guinan et al., 2003; Zhao & Dhar, 2009). Therefore, a key to understanding the contributions of the two reflexes is accurate detection and measurement. As a first step, here we propose an approach to MEMR detection based on a click stimulus time series analyzed in discrete frequency bands. No attempts were made in this work to study the effectiveness of this MEMR detection method in mitigating MEMR effects on the MOCR.

There have been multiple efforts among researchers to develop methods capable of detecting MEMR activation during MOCR measurements. Backus and Guinan (2007) introduced a stimulus frequency (SF)OAE phase gradient-based method of MEMR detection. The rationale was that MEMR-mediated changes in the SFOAE delay would be much shorter compared to MOCR-mediated changes. Other researchers have used concurrently presented tones (602 Hz and/or 226 Hz) to monitor the MEMR during SFOAE measurements (Goodman and Keefe, 2006; Zhao & Dhar, 2009). Abdala, Mishra, and Garinis (2013) monitored stimulus changes in their two-tone distortion product (DP)OAE stimulus to detect MEMR activation. The rationale for these latter two methods was that the MEMR activation would alter stimulus reflectance, causing a change in the ear canal stimulus pressure. Based on work by Feeney, Keefe, and Marryott (2003), Abdala et al. (2013) suggested that a stimulus level change of 0.12 dB (1.4%) is indicative of MEMR activation.

More recent investigations have employed transient stimuli such as clicks and tonebursts to detect the MEMR. Boothalingam and Purcell (2015) applied the same approach as Abdala et al. (2013) to clicks and reported no MEMR activations for low-level click/contralateral noise combination (55 dB peSPL/60 dB SPL, respectively). Mertes (2020) established critical differences in click stimulus for detecting MEMR based on group data. Marks and Siegel (2017) monitored tonebursts in a level-series. Goodman and colleagues (Goodman et al., 2013; Mertes and Goodman, 2015) used a resampling-based approach to statistically establish whether the changes in the stimulus level indicate MEMR activation. Goodman et al. (2018) included stimulus phase in the measurements, noting that while including phase increased the sensitivity of MEMR detections, it was not possible to disentangle the MEMR effects on the MOCR when both reflexes were active. Results from a majority of the above studies corroborate findings by Feeney and Keefe (Feeney et al., 2003; Feeney and Keefe, 2001) that the MEMR is activated at much lower levels (up to 21 dB) than that estimated by clinical instruments. Therefore, it is imperative that establishing the presence/absence of the MEMR in MOCR assays is not assumed based on MEMR thresholds from standard clinical instruments.

Despite the significant improvements made in recent years, uncertainty remains regarding how best to ascertain whether observed changes in OAEs represent purely MOCR-mediated activity or are a combination of both MEMR and MOCR activity. One reason is that MEMR effects are frequency dependent, resulting in increased stimulus reflection at the eardrum at lower frequencies but reduced reflection at higher frequencies (Feeney and Keefe, 1999; Borg, 1968). As such, when frequency-specific changes are not considered (e.g., Boothalingam and Purcell, 2015; Mertes, 2020), MEMR detection may be less sensitive. This is because MEMR-induced stimulus changes at the lower and higher frequencies may partially cancel out and falsely reflect little or no change. Such cancellations may be a larger issue for broadband stimuli such as clicks than for DPOAE- or SFOAE-based assays.

Another palpable reason for the uncertainty is that the presence of MEMR in all the aforementioned methods is deduced by comparing the change in stimulus magnitude from only two conditions: with- and without-an acoustic efferent activator (typically contralateral noise). While resampling methods can indicate whether a reliable difference in stimulus level exists between the two conditions, the difference cannot always be attributed to the MEMR. Stimulus pressure level changes due to probe movement and/or changes in middle ear pressure during recording can also produce stimulus level differences between conditions (Goodman et al., 2013). For this reason, evaluating the time course of the recorded stimulus level may be useful. The MEMR demonstrates an exponential onset that varies with the stimulus level (Hung and Dallos, 1972). This predictable behavior can be leveraged to identify true MEMR responses, both visually and statistically.

The purpose of the present study was to address these sources of uncertainty by introducing a method that evaluates stimulus pressure recordings of click trains in order to identify the presence of MEMR. The kinetics of the resulting time series are evaluated in a frequency-specific manner. When paired with statistical tests, the approach provides a sensitive, objective method for detecting middle ear pressure changes over time and, with further validation, will allow for discrimination between changes due to the MEMR versus other sources.

## II. METHOD

### II.A. Participants

Twenty young clinically-normal hearing volunteers (mean age: 22±2.7 years; 2 males) participated in the study. Clinically normal hearing was established by an unremarkable otoscopy examination, behavioral hearing thresholds ≤20 dB HL between 0.25 and 8 kHz (SmartAud, Intelligent Hearing Systems, FL), type-A tympanograms (Titan, Interacoustics, Denmark), and measurable distortion product (DP) OAEs (0.5-6 kHz; 1/3^rd^ octave intervals; 6 dB signal-to-noise ratio [SNR] criterion) (SmartDPOAE, Intelligent Hearing Systems, FL). Middle-ear muscle reflex thresholds were also measured in all participants using a clinical system (Titan, Interacoustics, Denmark; see also Section II.F). All participants provided written informed consent prior to participating in the study. Their participation counted towards extra-credit for various courses in the undergraduate program within the Department of Communication Sciences and Disorders. All study procedures were approved by the Health Sciences Institutional Review Board at the University of Wisconsin, Madison.

### II.B. Experimental set-up

Stimuli were digitally generated in Matlab (Mathworks, MA) using an iMac (Apple, CA). Synchronous playback and recording of signals were done using the Auditory Research Lab Audio Software suite (ARLas; Goodman, 2017) in Matlab. The iMac was interfaced with a sound card (Fireface UFX+, RME, Germany) for analog-to-digital-to-analog conversion at a sampling rate of 96 kHz and bilateral stimulus delivery to the participants’ ears was achieved using a dual-probe ER10X system (Etymotic Research, IL). The ER10X probe microphones recorded the ear canal pressure bilaterally. The ER10X probes are heavier than other Etymotic Research probes (e.g., ER10B+ and ER10C) and use plastic as opposed to foam tips to couple the probe to the ear. As a result, the probes tend to slip out of the ear during long recording periods. To avoid such probe slippage the ER10X probes were held securely in place in participants’ ears by (1) hanging the cables from the sound booth ceiling, (2) attaching them to hollowed-out earmuffs with only the headband and cushion in place (Mpow 035, Mpow, CA), and (3) puttying around the probes in-ear using silicone earmold putty (Silicast, Westone Laboratories, CO). In addition, an in-situ stimulus calibration was done before the start of every measurement condition. The calibration results, forward (FPL) and sound pressure (SPL) levels for a broadband chirp (0.2-20 kHz), were monitored throughout the experiment for gross deviations in spectral level. Along with the temperature and humidity sensors and the heating element in place in the ER10X probe, to avoid drifts in microphone sensitivity, the measures taken in the study successfully mitigated deviations in the stimulus spectral level over the course of the experiment across all participants.

The experiment was conducted in a double-walled sound-attenuating booth where participants sat comfortably in a recliner for the duration of the experiment and watched a silent closed-captioned movie. Participants were asked to sit relaxed, not move, and swallow as few times as comfortable, and not sleep. They were allowed to move and do any noisy activities (e.g., cough) between each experimental condition, i.e., roughly every 10 mins. The experimental procedure took about an hour to complete with about 1-2 min breaks between conditions. Consenting and screening procedures took an additional 30-40 mins.

### II.C. Stimulus, calibration, and paradigm

The level of the clicks was varied from 65 dB peak-to-peak (pp) SPL to 95 dB ppSPL in 6 dB steps in 6 separate stimulus level *conditions*. We refer to ppSPL as the SPL generated for a given peak-to-peak pressure fluctuation within the stimulus, i.e., without referencing the peak-to-peak of the click to the peak-to-peak of a 1 kHz tone. The latter is typically referred to as the peak-to-peak equivalent (pe) SPL (Burkhard, 2006). Stimuli were presented bilaterally. Each *block* consisted of a 1.008 s long click train, (henceforth referred nominally as 1 s) and a 0.742 s silent period. Each block was repeated 335 times per stimulus level condition. The click train consisted of 63 clicks presented at a rate of 62.5 clicks per second. This click rate translated to an *epoch* duration of 16 ms. Unlike a traditional contralateral noise elicitor paradigm, we used the OAE-evoking clicks to also elicit the MEMR. Prior studies have demonstrated that clicks indeed elicit the MEMR with a monotonic reduction in the reflex threshold with increasing click rate from 50-200 Hz (Rawool, 1995;1996; Johnsen and Terkildsen, 1980). While a faster (>62.5 Hz) rate may elicit more robust MEMR, the goal of this study was to detect the MEMR during MOCR measurements. The 62.5 Hz rate used in this study was conducive for this purpose because it elicits robust MOCR at 80 dB ppSPL (Boothalingam et al., 2018) while allowing for the recovery of OAEs in the 16 ms epoch duration. Any faster rate would not allow for OAE recovery in the frequencies of interest (1-4 kHz). The 1 s long click train was expected to allow the MEMR to reach steady state (Hung and Dallos, 1972).

Click stimuli were generated in the frequency domain. Click spectra were bandlimited between 0.8 and 6 kHz and flattened at the eardrum using FPL calibration, in order to homogenize cochlear stimulation across participants by accounting for the differences in external and middle ear impedance characteristics. Detailed descriptions of FPL calibration can be found in other reports (e.g., Scheperle, Neely, Kopun and Gorga, 2008; Souza, Dhar, Neely, and Siegel, 2014; Dewey and Dhar, 2017). Briefly, prior to data collection, Thevenin-equivalent source calibration of the probes were obtained for known acoustic loads of tube lengths 2.9, 3.6, 4.15, 5.2, 6.9 cm (diameter = 8 mm). The different lengths of the tube were achieved by moving the piston in the inbuilt calibration cavity of the ER10X system. The load (participants’ ear) calibration was performed before each condition, i.e., roughly every 10 mins, to obtain the ear canal and middle ear impedance, surge impedance, and the pressure reflectance. These estimates were used to calculate the forward and reverse pressures in the ear canal and build the external and middle ear transfer function (Rasetshwane and Neely, 2011). Based on the transfer function between 0.8 and 6 kHz, correction factors were created to generate a flat spectrum click at the eardrum. This matrix was then inverse Fourier transformed to the time domain to make the click. Clicks were then scaled to obtain the desired in-ear ppSPL level.

Pilot work suggested that in some participants, it was not possible to achieve the desired level at the highest click level condition (95 dB ppSPL) due to loudspeaker output limitations. To avoid this issue and reduce loudspeaker ringing, all the aforementioned calibration procedures, including source calibration, were done for clicks simultaneously presented through the two loudspeakers of each probe. This method allowed the highest click level to be reached by increasing the level of the click by 6 dB. The duration of the final acoustic click and its ringing in an ear simulator was roughly 3.5 ms.

### II.D. Response analysis

Recorded ear canal stimulus pressure was first bandpass filtered between 0.7 and 6 kHz. Although the click stimulus bandwidth extended to 6 kHz, preliminary data analysis and prior studies have suggested minimal MOCR activations past 3.5 kHz (Goodman et al., 2013). Therefore, the click stimuli were considered only between 0.89 to 3.5 kHz in the analyses reported below. After filtering, post-hoc artifact rejection was applied.

Epochs with root-mean-squared (RMS) amplitude >2.25 times the interquartile range were discarded as artifacts (Goodman, Fitzpatrick, Ellison, Jesteadt, and Keefe, 2009). The rejection rate by this method was approximately 10% of recordings, across participants.

#### II.D.1. Stimulus extraction

Stimulus level was estimated from ear canal stimulus pressure waveforms that were time windowed between −1 and 3.5 ms, with time zero defined as the location of the maximum excursion of the click pressure waveform. Raised cosine onset and offset ramps (0.5 ms duration) were applied to the windowed pressures. Waveforms within this time window were converted to the frequency domain via FFT, and the resulting SPL was separated into FPL and reverse pressure level (RPL) components using the source and load impedances obtained during stimulus calibration (Rasetshwane and Neely, 2011; Scheperle et al., 2008). We observed that using RPL, FPL, or SPL estimates of the stimulus level did not produce any difference in the results. This may be because our measure of FPL included both the incident pressure and all subsequent forward-going energy. Similarly, RPL included all reverse going energy. We chose to use RPL in all further analyses.

#### II.D.2. MEMR estimation

The RPL vector (complex vectors in the frequency domain) were arranged into a 3-dimensional (3D) matrix, X, with *i* = 2674 rows (Fourier frequencies), *j* = 63 columns (points in the time series) and *k* = 335 repetitions. That is, there were *k* repetitions of each row and column (*i,j*). The noise floor at each frequency and time point was calculated as the standard error of the mean across the *k* repetitions (Goodman et al., 2009). The magnitude at each frequency and time point was calculated as the magnitude of the mean across the *k* repetitions. The magnitudes were then reduced from the full set of *i*=336 Fourier coefficients to to *i*=6 values corresponding to the passband frequencies of a bank of third-octave filters with center frequencies *f*= 1, 1.3, 1.6, 2, 2.5, and 3.2 kHz. The reduction was accomplished by taking the energy-weighted average within each frequency band and was performed separately for each point in the time series. Additionally, synchronous spontaneous OAEs (SSOAEs) were removed at this time, as described in section II.E. This reduction process reduced the original data matrix to a 6×63 two-dimensional matrix, i.e., the time series for each of the six frequency bands. For each frequency band, the magnitude in Pascals at each point in the time series was divided by the magnitude in Pascals at the first time point. This metric of relative change, referred to hereafter as Δ, was then expressed in dB. This operation is equivalent to subtracting the magnitude in dB SPL at each time point with the magnitude at the first time point in dB SPL.

At each of the six frequency bands, relative magnitude (Δ) across the 63 points in the time series provided an estimate of the change in the stimulus magnitude over the course of 1 s. No change in the stimulus corresponded to 0 dB, while positive or negative values were interpreted as an increase or decrease in the stimulus magnitude relative to the magnitude at the first time-point. The MEMR time series from the majority of participants displayed monotonic exponential change over time. A typical example is shown in Fig. 1A. A smaller subset of participants (4 out of 20) showed more complex, non-monotonic change in some frequency bands. A representative example is shown in Fig. 1B. Two differences between the data in panels Fig.1A and B can be appreciated: (1) the change in Δ, especially at 1 kHz, occurs in opposite directions and (2) for the same 1 kHz band, there is a momentary reduction in magnitude before it increases. The magnitude and time-constants of the MEMR were estimated from the time series using a physiologically motivated exponential-model fit to the data, similar to those used for the MOCR (Liberman et al., 1996; Kim et al., 2001; Meinke et al., 2005; Backus and Guinan, 2006; Srinivasan et al., 2012). Like the MOCR, the time series of the MEMR also displayed a *latency* in its onset in majority of the participants (Backus and Guinan, 2006). Exponential fits, one- or two-term, did not model this behavior adequately. Therefore, we visually inspected the Δ at the 95 dB ppSPL condition to mark the onset of stimulus change, i.e., a marked departure from zero change. The 95 dB ppSPL condition was used for this purpose to identify this apparent latency with the highest SNR and the largest response possible. The Δ values prior to the onset were excluded from the fitting process. More details about the onset latency are provided in the Results section (see Section III.C). We found that a single-term exponential did not adequately describe the data even when only the data after the onset were considered, but a two-term exponential did. We therefore fit each time series with a curve of the form:

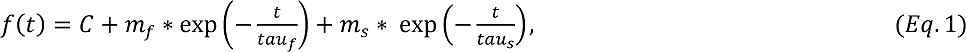

where, *f(t)* is the fit as a function of time *(t)*. The variables *m_f_* and *m_s_* are the magnitudes of the fast and slow components of the fits, respectively. The variables *tau_f_* and *tau_s_* are the fast and slow time-constants, respectively. *C* is a constant term representing offset along the y-axis. The estimate of MEMR magnitude change, hereafter referred to as **MEM**, was the final value (at time 1 s) of the fitted function. This value represented the magnitude of change after 1 second, relative to the starting value. Minimum and maximum bounds for the two time constants were set based on prior work (Dallos, 1964; Hung and Dallos, 1972). The *tau_s_* was allowed to range between 1 and 500 ms and the *tau_f_* was allowed to range between 1 ms and 50 seconds.

**Fig. 1.**
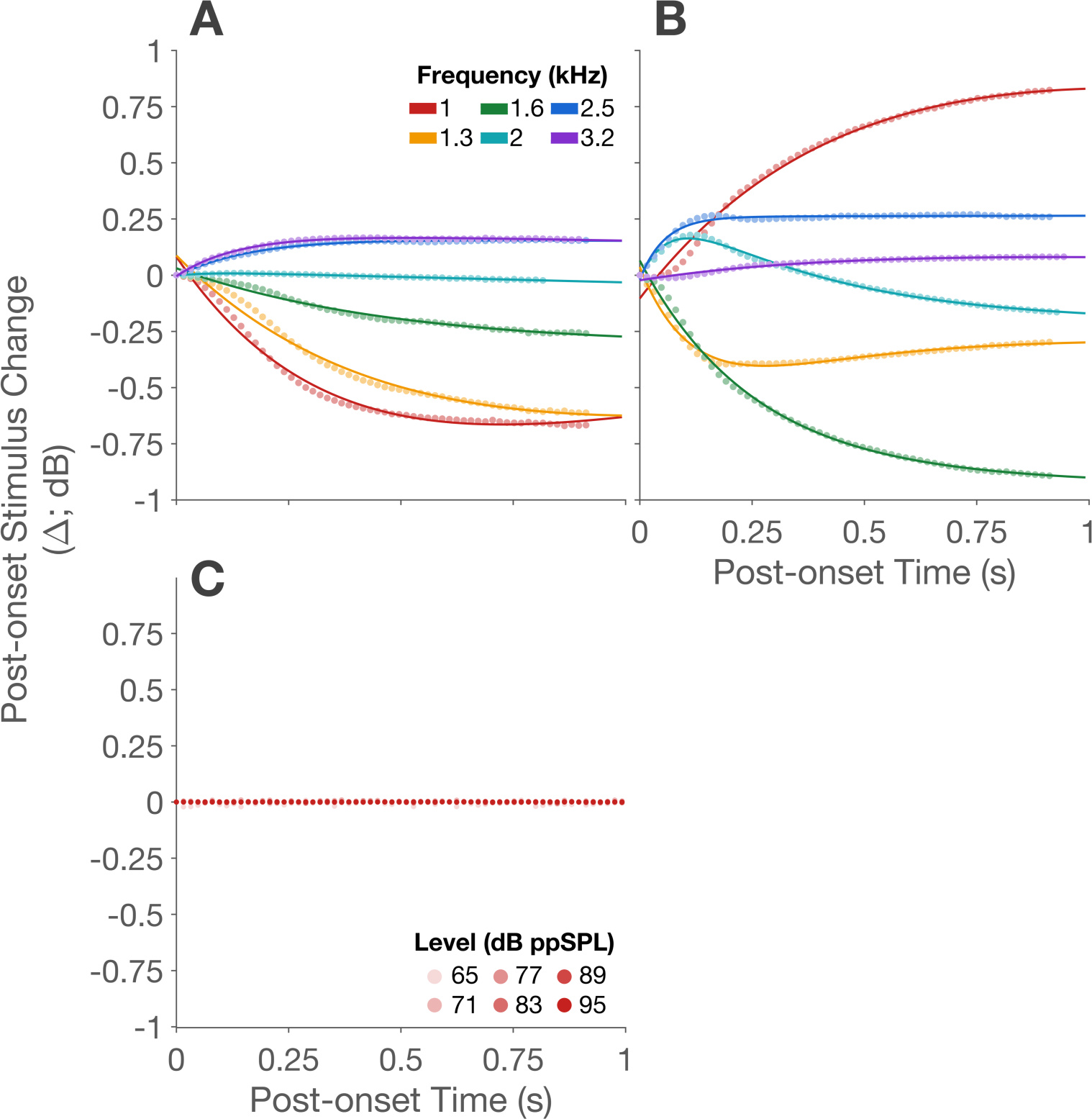
(color online) Panel A shows one example of typical, monotonic, MEMR time course and Panel B shows one example of non-monotonic MEMR time course from two different participants. In both top panels, colors indicate the third-octave band center frequency and scatter points are the change in stimulus level post onset re: the first time point. The line fits are two-term exponential models fit to the data (see Eq. 4). Data from the experiment run in an ear simulator is shown in Panel C. Only the 1 kHz data, but at all stimulus levels, are shown for clarity.

### II.E. Measures to avoid MOCR influence on MEMR measurements

OAEs may contaminate the stimulus pressure estimation if they are included in the stimulus window. This could be a problem because MOCR-mediated changes in the OAE could masquerade as MEMR activity. The restricted frequency range of 0.89-3.5 kHz in the study was expected to avoid contamination from click-evoked (CE) OAEs in the stimulus window. Given the dispersion in OAE frequency-latency relationship of reflection-generated emissions (e.g. SFOAEs and CEOAEs), it is highly unlikely that emissions below 3.5 kHz would be included in the stimulus window (<3.5 ms; Shera, Guinan, and Oxenham, 2002). Even if some emission pressure was included in the stimulus window at higher stimulus levels, at these high levels the stimulus sound pressure would dominate and any emission pressure would likely add an insignificant amount to the stimulus. Any change in CEOAEs due to the MOCR is also unlikely to be significant enough to influence the MEMR measurements (Keefe, Fitzpatrick, Liu, Sanford, and Gorga, 2010).

Spontaneous and synchronized spontaneous (SOAEs and SSOAEs) may pose a greater challenge. These emissions are long lasting and can be present in the stimulus window. To avoid these influences, we removed frequencies containing S/SSOAEs from the click stimuli. The last click epoch of the 95 dB ppSPL condition was used to monitor S/SSOAEs in a temporal window extending from 20-40 ms after the click presentation. Ten such windows were concatenated before performing an FFT, to improve the spectral resolution. The resulting FFTs were averaged and windowed in the same frequency range as the click stimulus (i.e., 0.8-3.5 kHz). S/SSOAEs were identified as spectral peaks ≥10 dB above the noise-floor (similar to Marshall, Miller, Guinan, et al., 2014; Lewis, 2020). Of the 20 participants in the study, 12 (60%) had 56 measurable S/SSOAEs in total. The mean S/SSOAE magnitude was −9.7 dB SPL with magnitudes ranging from −23.4 to 9.2 dB SPL. The median number of S/SSOAEs per participant was 1.5 with a maximum of 9 in three of the twelve participants. The prevalence of S/SSOAEs in the present study is smaller than prior some reports (72% to 85%; Keefe, 2012; Sisto, Moleti, and Lucertini, 2001). However, it is within the range for SOAEs which range from 50% to 70% (Talmadge, Long, Murphy, et al. 1993; Abdala, Luo, and Shera, 2017).

Following S/SSOAE identification, contaminated FFT frequency bins were assigned a weighting of zero in the energy weighted averaging processes described above. There are some potential limitations to this method of S/SSOAE detection and removal. S/SSOAEs outside of the frequency range of CEOAE measurement have been reported to influence CEOAEs (Keefe, 2012). Therefore, despite removing S/SSOAEs within the frequency range of interest, S/SSOAEs outside this range may have influenced the stimulus. However, S/SSOAEs are most prevalent in the 1-4 kHz range (Abdala et al., 2017; Keefe, 2012; Jedrzejczak, Blinowska, Kochanek, et al, 2008; Talmage et al., 1993) where the current S/SSOAE analysis was conducted. Therefore, influence from S/SSOAEs outside the range we tested is likely to be small. Notwithstanding, we report the median and the interquartile range (25% and 75% percentile) values for MEM, tau_!_ and tau_%_ for analyses with S/SSOAEs removed and S/SSOAEs included. Overall, the raw values, and visual and statistical re-examination did not reveal any differences for MEM. Minor differences (on the order of 4%) were seen for tau. To be conservative and avoid confusion, only the statistics for data with S/SSOAEs removed are presented. This lack of qualitative and quantitative difference between the time series data and overall results with and without S/SSOAEs suggests that the frequencies outside this range did not substantially affect stimulus magnitude.

### II.F. Comparison with a clinical instrument

MEMR elicited with broadband noise in a clinical instrument (Titan, Interacoustics, Denmark) was also measured for purposes of comparison with the MEMR threshold estimated using the proposed method. While not the focus of the study, this comparison allowed for corroboration of lower thresholds in click-based wideband MEMR estimation relative to clinical instruments reported by others (e.g., Feeney and Keefe, 2001; Feeney et al., 2003; Marks and Siegel, 2017). To be consistent with the studies in the literature, we used the 226 Hz probe tone to monitor changes in magnitude of admittance. Threshold from the clinical tympanometer was calculated as the lowest level at which the activator (broadband noise) caused a 0.02 ml change in admittance.

### II.G. False positive check

To ensure the changes in stimulus levels were unrelated to system artifacts or other non-physiological factors, we ran all conditions of the experiment in an ear simulator (Type 4137; Bruel & Kjaer, Naerum, Denmark). The outputs from the analysis of this data for the 1 kHz band across levels are plotted in the lower panel of Fig. 1. There was no evidence of any systematic changes in the stimulus estimate across time and level. No line fits are shown because none of the fits were significant (explained below). Compared to normal hearing participants, the range of random stimulus changes in the ear simulator was 2-3 orders of magnitude smaller. We also did not find any frequency effects (data not shown for ease of visualization). This check provides confidence that any changes in the stimulus that we observe must have physiological origins and were not system related.

### II.H. Statistical tests

#### II.H.1. Significance test of fits to data

MEMR activity was considered possible at a given frequency and level if 1) the time series showed systematic (non-random) change across time and 2) the systematic change was also present in that same frequency band at all higher stimulus levels. The presence of systematic change was tested using the Heller-Heller-Gorfine test (HHG; Heller, Heller, and Gorfine, 2013). The HHG was designed as a nonparametric test of association between two random vectors. It is implemented between two vectors of the same dimensions and is sensitive to nearly any form of dependence. If both of the vectors are composed of non-random sequences, the test will report an association. We implemented the test by letting the first vector be a possibly random MEMR time series sequence. The comparison vector was the two-term exponential fit to that same sequence. Implemented in this manner, we tested the null hypothesis that there was no association between the two vectors, i.e., that both vectors were random sequences.

Since the vector representing the exponential fit was never a random sequence, we were in effect testing whether the MEMR time series showed any systematic change over time. If the MEMR time series showed any systematic change across time, the HHG test detected a dependence between the two vectors, the null hypothesis was rejected, and that time series was considered to show evidence of either MEMR, some other systematic change over time, or both. Significance of the comparison (*p*-value) was obtained by generating confidence intervals from bootstrapping the HHG test 1000 times. Because each participant had data at six frequencies at each of the six stimulus levels, *p*-values were corrected for alpha inflation using the False Discovery Rate (FDR; Benjamini and Hochberg, 1995) method.

If a systematic change detected by the HHG test was not persistent at all higher stimulus levels, then MEMR activity was not considered to have occurred at that level. For each individual, MEMR threshold was considered the lowest stimulus level at which at least one of the six frequency bands showed significant MEMR activity as just described.

#### II.H.2. Group statistics

To study MEM and tau group data, a linear mixed-effects model (*lme4* package in R; Bates, Mächler, Bolker, and Walker, 2015; R Core Team, 2014) was used. Dependent variables MEM and tau were fit with (1) *fixed-effects* of predictors level (*L*), frequency (*f*), and the interaction between level and frequency (*L*X*f*) and (2) *random-effects* of participants and random slopes for level. Stimulus level was treated as a continuous variable because levels were incremented in discrete steps and frequency was treated as a categorical variable because Δ was discretized by averaging in 1/3^rd^ octave bands:

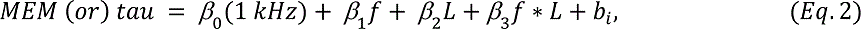

*β_0_* is the intercept when level is at the origin and frequency is 1 kHz, *β_1_* for each of the five frequencies (1.3, 1.6, 2, 2.5, and 3.2 kHz) is the simple effect, i.e., the intercept, of frequency when the level at origin with 1 kHz as the reference condition because frequency is treated as a categorical variable. *β_2_* is the slope of the level, *β_3_* is the interaction of level and frequency, i.e., the difference in the simple slopes of level for the six frequencies, and *b_i_* is the random-effects intercept for each participant *i*. The model was tested for significance using analysis of variance (ANOVA).

To study MEMR detections, percentage detections, hereafter referred to simply as detections, were calculated as number of participants with significant MEM for each frequency divided by the sample size (20). A general linear mixed-effects model was fit to the dependent variable detection in a fashion similar to the linear mixed-effects model for MEM and tau (Eq 4), except logistic regression was used instead of linear regression as the data were binominal (MEMR present vs. absent as determined by the HHG test). The log-odds of MEMR presence (logit) were obtained from the model. A logit link function, f(μ) = log(μ⁄(1 − μ)), where μ is the mean of MEMR presence, was used to transform the probabilities to a continuous percentage scale.

## III. RESULTS

### III.A. Stimulus change in the time series shows frequency-specific changes in middle ear impedance

The increase in stiffness in the middle ear due to the MEMR action will generally attenuate low frequency energy from reaching the cochlea more than high frequency energy. However, this relationship is non-monotonic. Data from Feeney and Keefe (1999; 2001) on the change in wideband reflectance due to MEMR activation show a reversal, i.e., reduced reflectance, or increased absorbance, between 1 and 1.5 kHz. Based on these prior data, we expected a reduction in the recorded ear canal stimulus level for the lower third-octave bands (1-1.6 kHz) and a relative increase in the recorded ear canal stimulus level for frequency bands ≥1.6 kHz. The time series for the six third-octave bands for each stimulus level, averaged across participants, is shown in Fig. 2.

**Fig. 2.**
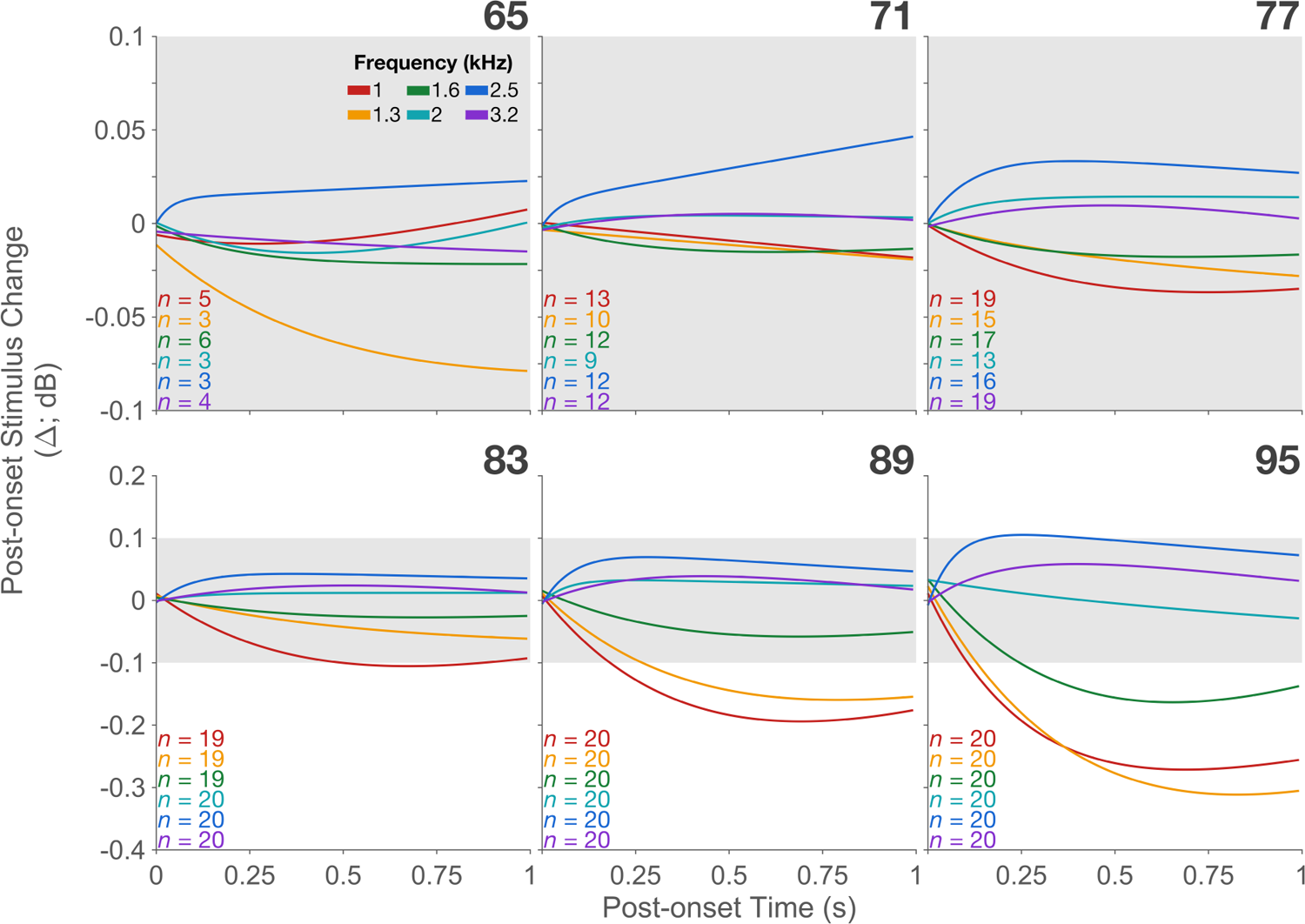
(color online) Group mean time series (panels by stimulus level). In each panel, colors represent the different third-octave frequency bands. Data are change in stimulus level post onset re: the first time point. Only the fits to the group mean data are shown for easy visualization. The y-axis of the top panels has been zoomed in to show smaller changes clearly. To maintain perspective, this zoomed region has been shaded with the same grey in the lower panels where the MEM is much larger. The n-size for each fit is provided within panels.

The time series of the MEMR at different frequencies was consistent with these expectations. The transition in the direction of change between the low vs. high frequencies appears to occur between 1.6 and 2 kHz corroborating prior reports (Feeney and Keefe, 1999; 2001; Schairer et al., 2007). An unexpected observation in the time series is that lower frequencies (1 and 1.2 kHz) appear to approach steady-state gradually close to 1 s, but the higher frequencies reverse course towards baseline shortly after onset, i.e., adapt to the presence of the stimulus.

The distribution of MEM across individuals is shown by the box plots in Fig. 3. MEM fits that did not pass the inclusion criteria (see Section II.H.1) were excluded from these plots. The number of participants at each level and frequency combination (*n*-sizes) are the same as those provided in Fig. 2. Level and frequency effects are apparent: MEM clusters around zero at lower levels and demonstrates a larger spread as level increases. When data from all frequencies and levels were pooled, the range (25% and 75% percentiles) of MEM was between −0.07 dB and 0.021 dB with the median at −0.0083 dB (S/SSOAEs removed). These values remained virtually unchanged when S/SSOAEs were included with the range between −0.07 dB and 0.022 dB and with a median of −0.0081 dB.

**Fig. 3.**
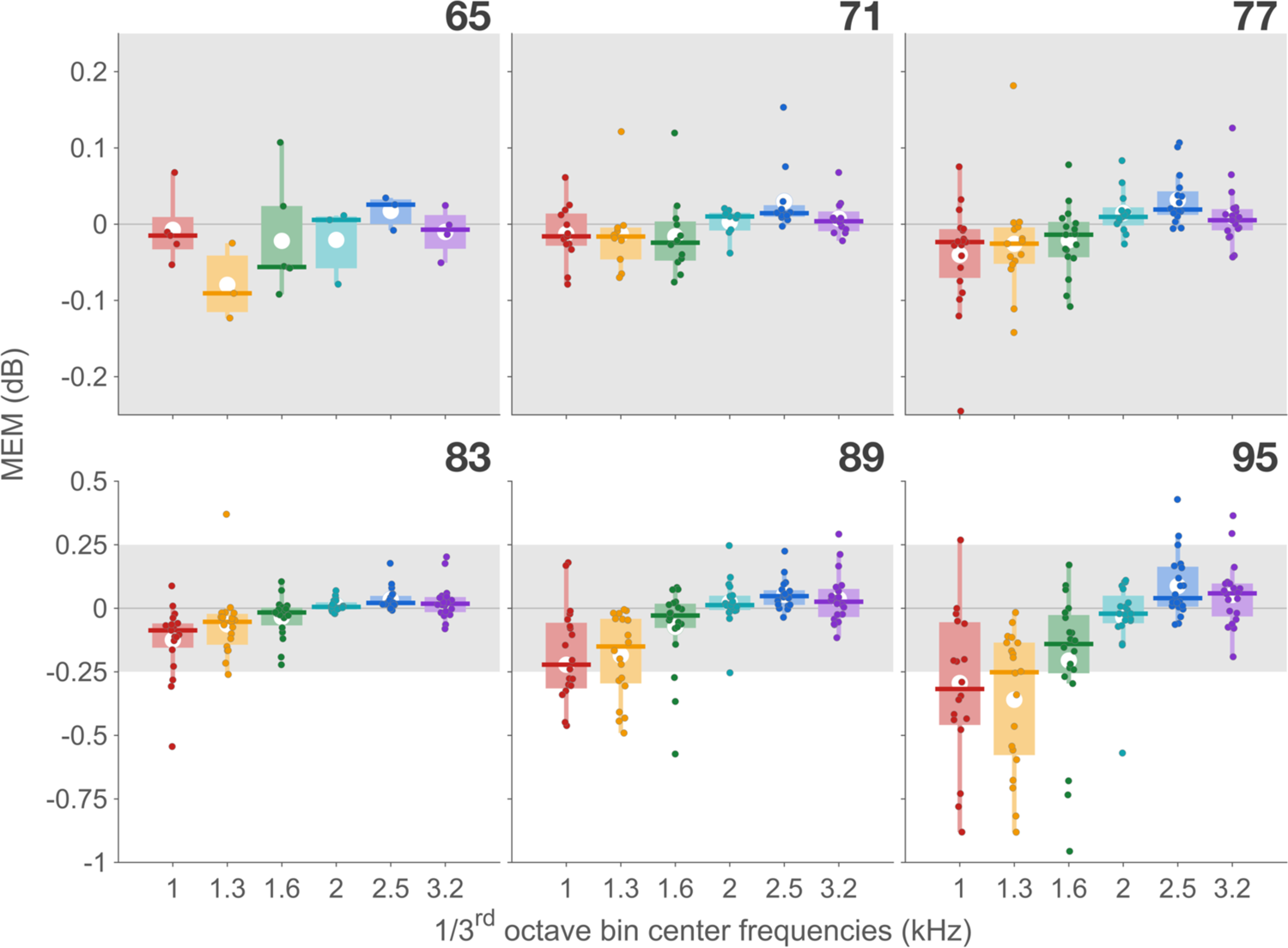
(color online) Box plots (panels by stimulus level) show individual MEM as filled colored circles on each box. Colors represent frequency in the x-axis. In the box plots, the white circle is the mean, and the vertical line is the data range. The horizontal, colored, line is the median. The y-axis of the top panels has been zoomed in to show smaller changes clearly. To maintain perspective, this zoomed region has been shaded with the same grey in the lower panels where the MEM is much larger.

### III.B. MEMR magnitude increases with level and decreases with frequency

In general, the data suggest that the Δ is larger at lower (<2 kHz) relative to higher frequencies (>2 kHz) at all stimulus levels. As seen in Fig. 4, the time series grew monotonically with increasing level at all frequencies. However, at higher frequencies (≥2 kHz), the time series tended to return to the baseline more strongly at the highest level. Note that these time series are averaged across participants where the time series for the same frequency can in some instances go in opposite directions (e.g., Fig 2B). The true magnitude of the MEMR can be compared across levels and frequencies when the absolute values (|MEM|) are plotted as input-output (IO) plots. In Fig. 5, IO for each frequency is plotted first in a separate panel and then summarized together in the top right panel (MEM-IO panel).

**Fig. 4.**
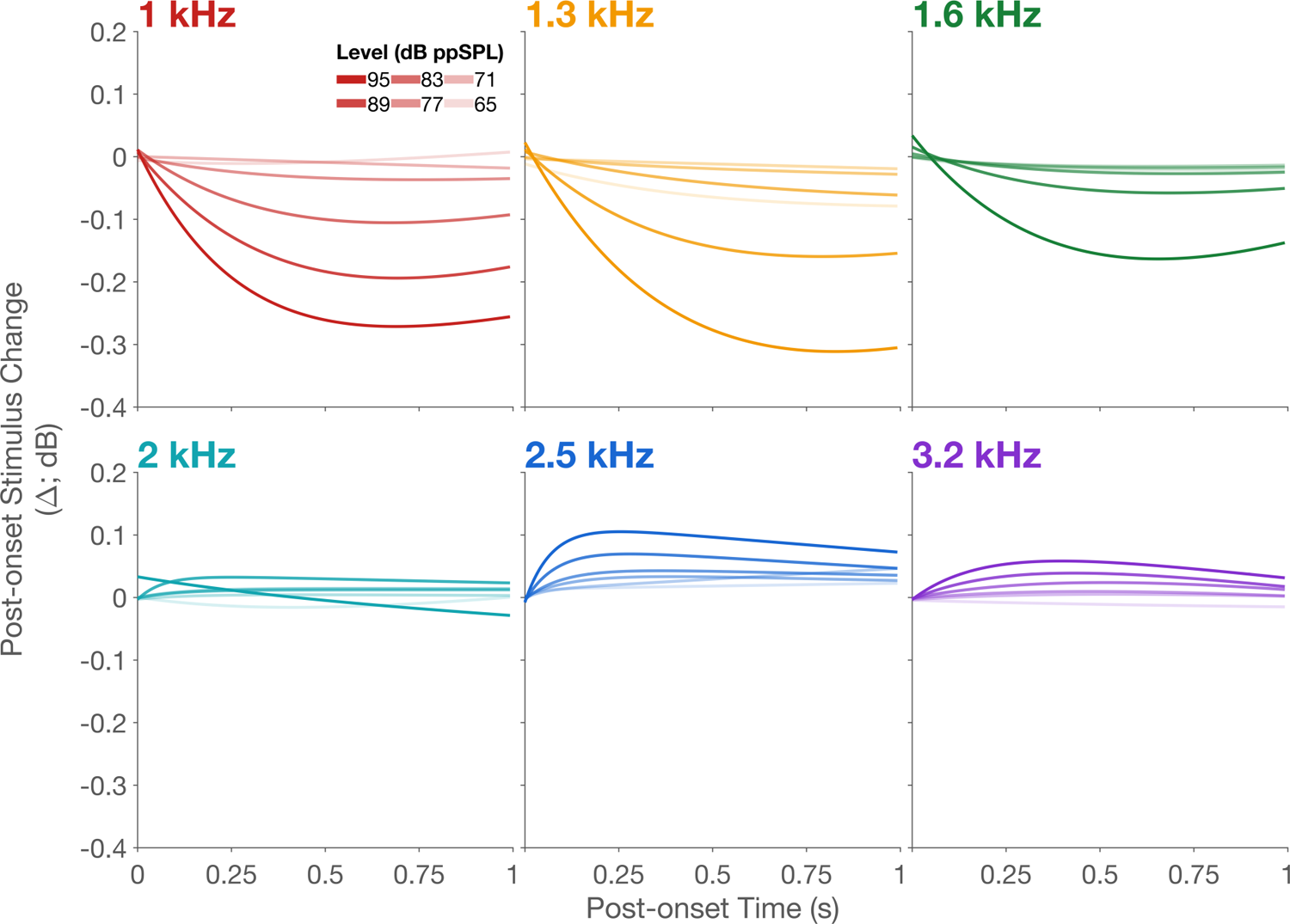
(color online) Group mean time series (panels by frequency). Data are change in stimulus level post onset re: the first time point. In each panel growth in MEM with increasing level, i.e., level series, is plotted. Darker the shade of a given color, higher the stimulus level. As expected, the magnitude of the time series increases with increasing level. Note the dichotomy in the direction of stimulus change as a function of frequency.

**Fig. 5.**
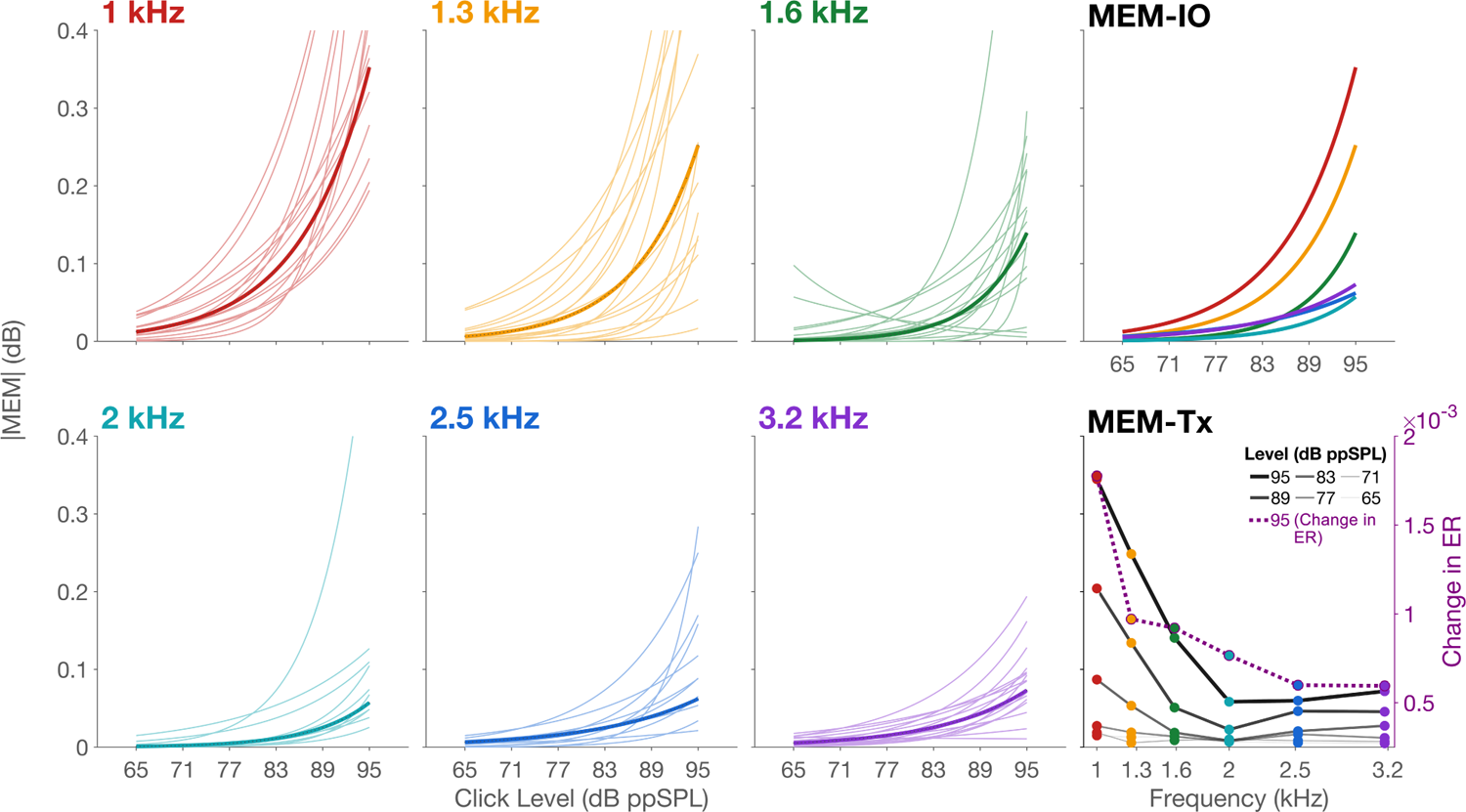
(color online) Input/output functions of the MEM are plotted in the left 6 panels (panels by frequency). Thin colored lines are individual one-term exponential fits to the absolute MEM at each level in a given frequency. The thick colored line in each panel is the one-term exponential fit to the mean MEM at each frequency. For a better comparison of this growth across frequencies, only the fits to the group mean are plotted in the panel MEM-IO. In the panel MEM-Tx, each line is a stimulus level with the MEM plotted as a function of frequency. This provides a transfer function of the MEM. Change in ER (only for the 95 dB ppSPL condition) is plotted on the right y-axis with data connected using a dotted line and circles outlined with purple color.

Growth of |MEM| was best approximated by a single exponential function. When the mean fits (thick lines) are viewed together (Fig. 5 MEM-IO panel) the differences in MEM growth with level as a function of frequency is apparent: lower frequencies undergo larger changes than higher frequencies. When the data are visualized as a function of frequency (Fig. 5 bottom right panel, MEM-Tx) with separate lines for each level, a MEM transfer function emerges. MEM grows non-monotonically with frequency and displays a minimum at around 2 kHz, most prominent at 89 dB ppSPL. To allow more direct comparisons with the data of Feeney and Keefe (1999), we also calculated energy reflectance, *ER*, as the squared magnitude of reflected pressure divided by the squared magnitude of the forward pressure. The change in ER time series was calculated in the same manner as MEM. Overall, the pattern of results for change in ER and MEM were similar. However, for reasons that are unclear, our mean ER change was two orders of magnitude smaller than those reported by Feeney and Keefe (1999; *n* = 3). When data are pooled across frequencies and levels, maximum and minimum ER were 0.0047 at 1 kHz and −0.00036 at 2 kHz, respectively. The change in ER for 95 dB ppSPL clicks are plotted in the MEM-Tx panel on the right y-axis.

To study the data inferentially, a linear mixed-effects model was used (see Section II.H.2). Only levels 77 through 95 dB ppSPL were included in the model as the lower levels had few MEMR activations. Given the exponential relationship between MEM and stimulus level, the values of MEM were linearized (by taking the logarithm of the absolute value) prior to fitting with a mixed-effects model. This transformation was necessary because linear models assume a linear relationship between the dependent and the predictor variables. Finally, to make the model estimates easier to interpret, we moved the origin of the predictor (level) to 77 dB ppSPL by subtracting the lowest stimulus level (77 dB ppSPL) from all levels. We first compared two models: with and without the level by frequency interaction term (*Eq. 2*). The interaction term was not significant (*p*=0.29) and was subsequently dropped from the model. The summary of results of the model is tabulated in Table 1.

**TABLE I.** Summary of ANOVA of the mixed-effects model with the dependent variable, MEM/*tau_f_*/*tau_s_*, and predictors, frequency, level (77-95), df = degrees of freedom, asterisks indicate significant effects.

| | ANOVA for MEM | | ANOVA for $\tau_f$ | | ANOVA for $\tau_s$ | |
| --- | --- | --- | --- | --- | --- | --- |
| Fixed effect variables | $F$-statistic<br>[df] | $p$-value | $F$-statistic<br>[df] | $p$-value | $F$-statistic<br>[df] | $p$-value |
| Frequency<br>($\beta_1$) | 25.5<br>[5,430] | <0.001* | 2.7<br>[5,316] | 0.02* | 1.4 [5,383] | 0.2 |
| Level ($\beta_2$) | 148<br>[1,431] | <0.001* | 1.3<br>[1,319] | 0.25 | 3.2 [1,384] | 0.07 |
| Frequency<br>($\beta_1$) X Level<br>($\beta_2$) | 1.2<br>[5,425] | 0.29 | 3.2<br>[5,316] | 0.008* | 4.9 [5,383] | <0.001* |

The lack of significant frequency by level interaction suggests that despite the difference in the magnitude of MEMR change with level at each frequency, the rate of change of magnitude with level is not different across frequencies, at least between 77 and 95 dB ppSPL. That is, the fit lines in Fig. 5 MEM-IO (top right) panel are different in their intercepts but not slopes (note that the exponential slopes were linearized by taking the logarithm of |MEM|).

The mixed-effects model analysis was followed by pairwise comparisons between frequencies with pooled level data, corrected for multiple tests using FDR. With the exception of three comparisons, 1 vs. 1.3 (*p*=0.2), 1.6 vs. 3.2 (*p*=0.3) and 2.5 vs. 3.2 kHz (*p*=0.2), all other paired comparisons were significant (*p*<0.042) with majority of the comparisons significant at *p*<0.001. These comparisons suggest that MEMR magnitudes at different frequencies (statistically modeled as the intercepts) were significantly different from each other, except the aforementioned three frequency pairs.

### III.C. MEMR kinetics demonstrate frequency X level interaction effects

Onset latency estimated at 95 dB ppSPL is plotted as a box plot in Fig. 6A. Data across frequencies were compiled for this box plot, as there was no significant effect of frequency (*F*[5,114] = 1.8, *p*=0.12) for latency in a one-way repeated measures ANOVA. The 25% and 75% percentiles for onset latency were 65 to 96 ms, respectively, with a median of 80 ms. Very few fits, 4 different frequencies in 6 different participants, had a sub-50 ms latency. However, 25 fits across all frequencies in 15 participants had latencies above 100 ms with a maximum of 210 ms.

**Fig. 6.**
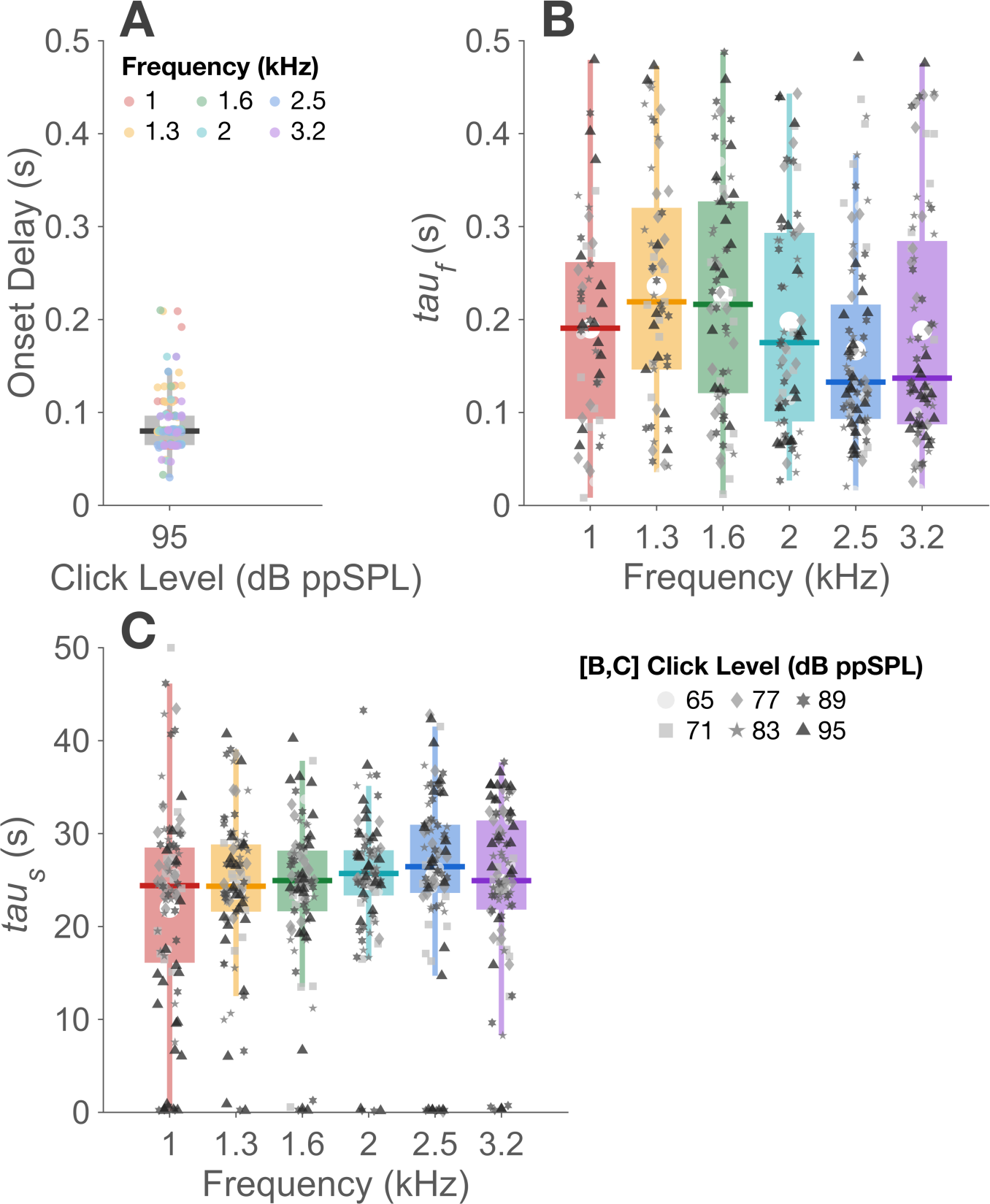
(color online) Box plots (panels by stimulus level) show MEMR kinetics. A. onset latency at 95 dB ppSPL with colored circles indicating the six frequency bands for all 20 participants (see Fig. 3 caption for details on box plots). B and C. *tau_f_* data and *tau_s_* data are presented, respectively, as box plots as function of frequency with pooled levels marked by varying symbols and shades of grey.

The distributions of the measure of kinetics, tau, are also plotted as box plots in Figs. 6B and 6C as a function of frequency. Instances where the *tau_f_* was greater than 0.49 s, bound set during the two-term exponential fitting process were not included. This rejection excluded 18% of the *tau_f_* data. The 25% and 75% percentiles for *tau_f_* were 98 and 281 ms, respectively, with the median at 175 ms when S/SSOAEs were removed.

Similar, to MEM, the *tau_s_* remained virtually unchanged when S/SSOAEs were included with range of 97.5 and 275 ms and a median of 168 ms. For *tau_s_*, especially at 89 and 95 dB ppSPL, the data contained several outliers with very small *tau_s_* values. To avoid outliers influencing statistical tests, we removed these outliers using the Tukey method where data points 2.25 times the interquartile range (IQR) were not included in the plot (Fig. 6C) and statistics. This process excluded 2.7% of the data points in *tau_s_*. The 25% and 75% percentiles for *tau_s_* were 21.8 and 29.3 seconds, respectively, with the median at 25.1 seconds with S/SSOAEs removed. With S/SSOAEs included, the range for *tau_s_* was 21.7 and 29.1 with the median of 25.1 seconds.

Similar to MEM (Fig. 3), only the levels 77 through 95 dB ppSPL were included in the linear mixed-effects model fit to both time constant data. The ANOVA results of this model for both time constants are presented in Table 1. Results from the mixed-effects model for the two time constants were slightly different. For *tau_f_* there was a significant effect of frequency but not for *tau_s_*. Level was not a significant predictor for either time constants (*p*>0.07). However, the level X frequency interaction was significant for both time constants.

Pairwise comparisons across frequencies corrected for multiple comparisons using FDR at each level also produced different results for the two time constants. For *tau_f_*, the interactions were driven mainly by the difference between 1 and 1.6 kHz in the low frequency range and 3.2 kHz in the high frequency range at all levels. For the 1 vs. 3.2 kHz, *p*=0.024, 0.025, 0.025, and 0.026 at 77, 83, 89, and 95 dB ppSPL, respectively.

For the 1.6 vs. 3.2 kHz, *p*=0.014, 0.016, 0.016, and 0.017 at 77, 83, 89, and 95 dB ppSPL, respectively. The mean differences were all positive in both cases, suggesting that 1 and 1.6 kHz had the longer time constants (re: 3.2 kHz). The results were reversed for *tau_s_*, where the interaction was mainly driven by the 1 kHz band but, here, the 1 kHz *tau_s_*, was smaller than other frequencies. The only significant comparisons were between 1 kHz and all other frequencies at all levels. All *p*-values were <0.033 with most values <0.02.

### III.D. MEMR detections increase with level

As shown in Fig. 7, and given the effect of level on MEM, it is not surprising that the number of participants identified as having significant MEMR activity in the group (detections) increased for all frequencies with increasing level (Fig. 7, bottom panels). This trend was best approximated by logistic fits with stimulus level as the predictor. All fits in this panel were significant (*p*<0.001).

**Fig. 7.**
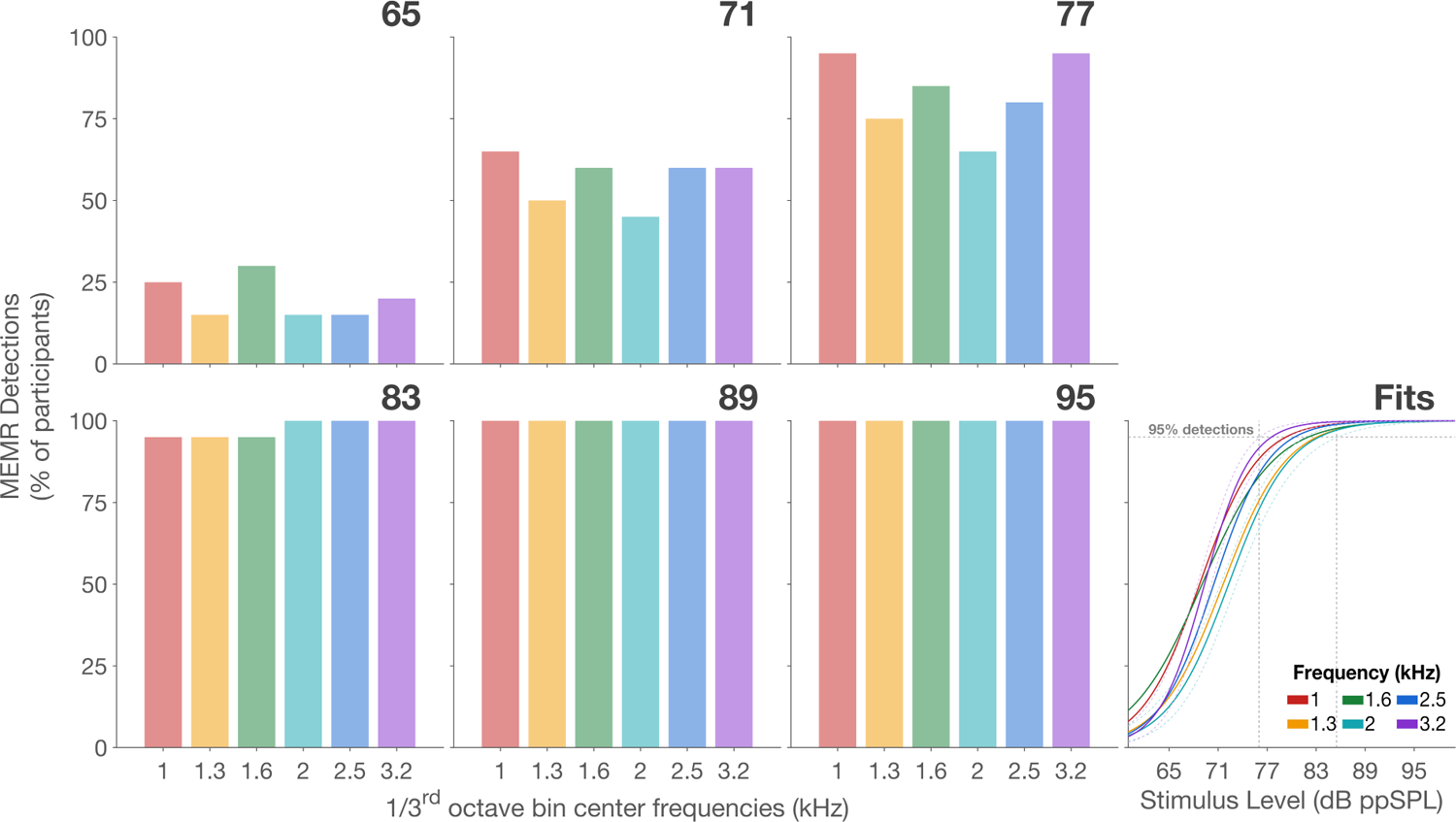
(color online) Percentage of MEMR detections as a function of frequency, at each level (separated by panels) is presented as bar graphs in the left 6 panels. In the Fits panel, these detections are fit with a logistic function with colors representing frequencies. Dashed lines around the steepest and the shallowest fits are included to show the 95% confidence of the fits. The horizontal dashed grey line indicates 95% detection. The vertical lines bookend the lowest and highest level, across frequencies, at which the 95% of the participants have MEMR activation.

To analyze the detections inferentially, we fitted data with the general linear mixed-effects logistic model (see Section II.H.2). Like the MEM mixed-effects model, the frequency by level interaction was not significant (*p*=0.16) and was subsequently dropped from the model. This new model results suggested that the level was a significant predictor of detections (*β_2_*; χ^2^ (1, *N*=20) = 138.1, *p*<0.001) but frequency was not (*β_1_*; χ^2^ (5, *N*=20) = 7.7, *p*=0.2). This result is consistent with the logistic fit-lines in the ‘Fits’ (bottom right) panel of Fig. 7. These results suggest that, unlike MEM, detections do not vary by frequency. By cautiously extrapolating these results to the population, it can be hypothesized that 95% of young normal-hearing individuals will show MEMR activation between 76 and 86 dB ppSPL at all frequencies (vertical dashed lines in Fig. 7 ‘Fits’ panel) when elicited using the stimulus and method proposed in this study.

### III.E. Comparison with clinical instrument

The MEMR thresholds obtained using the clinical instrument compared to those obtained using the study method are presented as a scatter plot in Fig. 8. Note that clinical MEMR threshold could not be obtained in one participant (technical issue) and in two participants the thresholds were higher than 95 dB SPL, the highest level tested. In the latter two instances, thresholds are shown as 100 dB SPL (open circles).

**Fig. 8.**
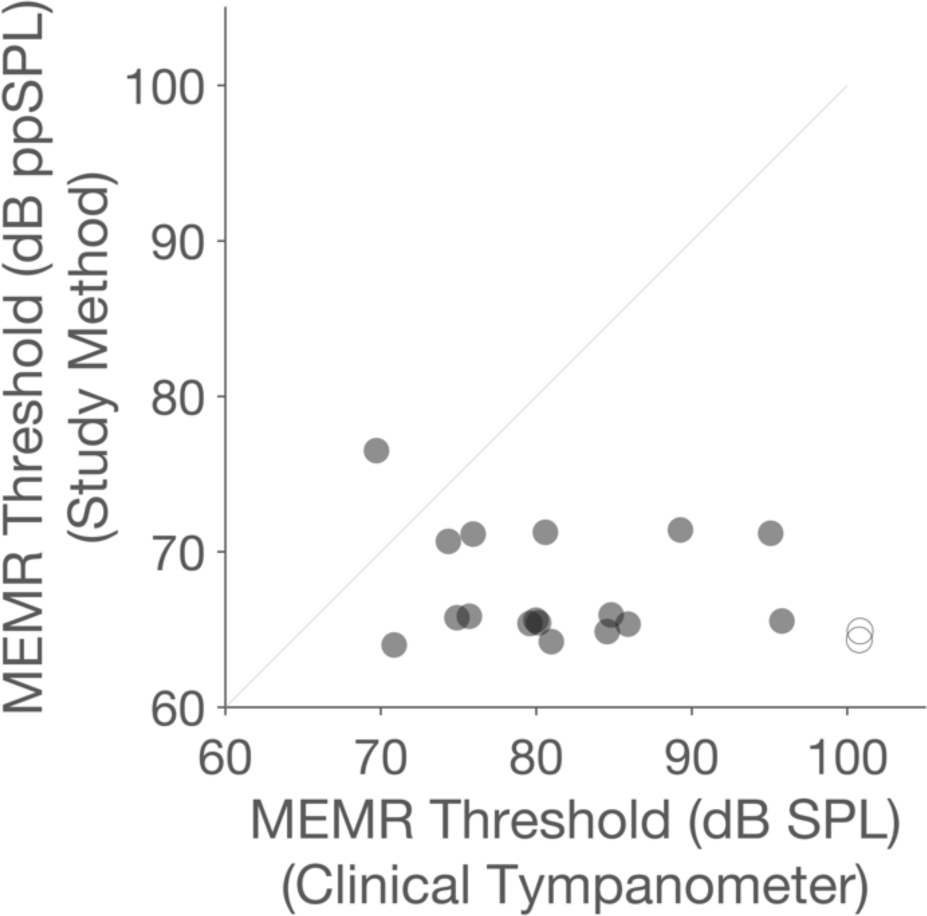
Scatter plot of MEMR thresholds obtained in the present study vs. those obtained for the same participants in a clinical tympanometer. The diagonal line shows unity relationship between the two variables.

Corroborating prior studies, the study method showed lower thresholds for all but one participant. The mean difference between the clinical method and the study method was 13.4 (SD = 8.9) dB, with the largest difference being 30 dB and the smallest difference −7 dB (1 participant), where the negative sign indicates the study method produced higher threshold. This result should be viewed cautiously due to differences in the stimulus type, level, mode of presentation (contralateral in clinical vs. bilateral in the study), atmospheric vs. tympanic peak pressure, and different criteria for defining a threshold in the study vs. clinical instrument. In the current study, threshold criterion was the presence of a statistically significant systematic magnitude change that was consistent at all higher stimulus levels at one or more of the six analyzed frequency bins. The number of frequencies that should demonstrate a significant and consistent change in order to most accurately determine MEMR threshold is currently unknown. Data from further studies should include individuals known to have no MEMR (e.g., individuals with profound hearing loss) to help establish threshold criteria.

## IV. DISCUSSION

The goal of this study was to describe a time series-based method of MEMR detection using a click train, with potential implications for OAE-based MOCR assays. Overall, our results showed that the click trains alone do elicit the MEMR and tracking the time series to detect MEMR is a viable method. Based on the agreement with frequency-specific changes in the MEMR (i.e., direction of change) reported from wideband estimates, the observed time series are consistent with increased stiffness in the middle ear system due to MEMR activation (Schairer et al., 2007; Feeney and Keefe, 1999; 2001).

### IV.A. MEM frequency effect is consistent with middle ear impedance changes due to increased stiffness

Given the direct relationship between resonant frequency (f) and stiffness reactance (X_s_) of a simple harmonic oscillator with a single stiffness and mass reactance (f ∝ X_s_), one would expect that stiffening the ossicular chain via MEMR activation would increase reflectance magnitude at the low frequencies and reduce reflectance magnitude at the higher frequencies. However, because the middle ear is a complex system with more than one mass and stiffness element, the relationship between impedance and frequency is non-monotonic. Specifically, Feeney and Keefe (1999), and subsequently further studies by Keefe, Feeney, and colleagues (Feeney and Keefe, 2001; Feeney et al., 2003; Schairer et al., 2007; Keefe et al., 2010) reported a prominent notch in the middle ear reflectance between 1 and 1.5 kHz, followed by a relative increase in reflectance beyond 2 kHz. The frequency effects observed in our data are consistent with the frequency-specific changes reported by Keefe and colleagues for MEMR-mediated changes in their click and chirp probes. When the frequency differences in admittance vs. reflectance are considered (see Fig. 6 of Feeney and Keefe, 1999) our results are also consistent with the results of Bennett and Weatherby (1979). As such, the stimulus changes observed in the time series in our data are consistent with the impedance changes engendered by MEMR activity.

The pattern of results across frequency we observed for MEM were similar to that reported for ER by Feeney and Keefe (1999). In that, the largest changes were seen at the 1 and 1.5 kHz bands with little change in the bands above 2 kHz (see Fig. 2 of Feeney and Keefe, 1999). However, the magnitude of the change in our *ER* calculations were smaller than those reported by Feeney and Keefe (1999) by two orders of magnitude. Other than the differences in the elicitors used in the two studies (tones in Feeney and Keefe, 1999), it is unclear what reasons must have led to these differences.

### IV.B. MEM frequency minimum matches impedance zero-crossing

In addition to the similarity in the general frequency effects observed between current data and prior work, the transition, or the zero-crossing, from decreasing to increasing reflectance between 1.6 and 2 kHz matches closely (Feeney and Keefe, 1999; 2001). Specifically, the minimum in the transfer function (MEM-Tx panel in Fig. 5) at 2 kHz, best observed at 89 dB ppSPL, illustrates this specific corroboration where the lowest values of MEM in the function are for the 2 kHz band. This 2 kHz transition frequency is observable in all of the wideband MEMR investigations for reflectance (e.g., Feeney and Keefe, 2001), admittance (e.g., Schairer, et al., 2007) and absorbed power level (Feeney et al., 2017). Based on the data of Keefe, Bulen, Arehart, and Burns, 1993, Feeney and Keefe (1999) argued that this null in impedance is due to the middle ear impedance being close to the characteristic impedance of the ear canal. As discussed below (see: Implications for MOCR assays), this null could be of particular importance for MOCR assays.

### IV.C. MEM adaptation is frequency-specific

An important observation in the current study was the difference in the shape of the time series between low and high frequencies (Fig. 4). With the exception of 1.6 kHz, it appears that the lower frequencies reach steady-state at around 500 ms and in general continue to stay in steady-state at the end of the 1 second block. However, the time series of frequencies 2 kHz and above tend to return towards baseline, adapting to the presence of the stimulus. This behavior may at first appear similar to the well-documented MEMR adaptation (for a review, see Wilson, Shanks, and Lilly, 1984).

However, this differential adaptation across frequencies is different from the MEMR monitored with a single low frequency probe (220 or 226 Hz) elicited by low- and high-frequency tonal elicitors. In our data, frequency-specific adaptation is seen in the wideband probe elicited by a wideband elicitor (click). Given that our probe is also the elicitor, the locus of this frequency-specific adaptation cannot be delineated into peripheral vs. central factors. However, there is abundant evidence that point towards a neural and/or central origin for frequency specific adaptation with tonal elicitors (Lutman and Martin, 1978; Mukerji, Windsor, and Lee, 2010; Wilson, Steckler, Jones, and Margolis, 1978). For example, the reflex decay test, which tracks MEMR adaptation, is commonly used in audiology clinics to delineate cochlear vs. retrocochlear pathologies, with faster adaptation for low frequency tonal elicitors taken as indications of auditory nerve or brainstem pathologies (Margolis and Levine, 1991; Stach, 1987; Wilson et al., 1984). We speculate that the adaptation seen in our data results from relaxation of some middle ear structure(s) (e.g., the stapedius muscle, tympanic membrane) over time resulting in a frequency specific change in impedance.

### IV.D. Idiosyncrasies in MEM time series likely reflect stapedius muscle behavior

Similar to the individual data shown in Fig. 1B, three other participants displayed non-monotonic time series, albeit with much slower transition from negative to positive (or vice versa) change in the time series. Although not captured by the fit adequately, a rapid alteration in direction of change exists at 1 kHz (Fig. 1B) and has been reported previously by Hung and Dallos (1972). These authors attributed such rapid transition in the direction of change to the stapedius muscle momentarily relaxing before contracting, in a phenomenon referred to as “latency relaxation”. Data from Hung and Dallos (1972) on the latency of this change in impedance are similar to the 100-150 ms seen in the present study. Without monitoring the MEMR time series, such idiosyncrasies will go unnoticed while producing puzzling and unexpected changes in reflectance or impedance. Therefore, there is value to monitoring the time course of the reflex in addition to estimating the final magnitude, in order to garner a better understanding of the underlying physiology.

### IV.E. MEM and detections increase with stimulus level

The MEMR estimate, MEM, was expected to grow with increasing level. This growth function, as seen in Figs. 5 and 6, is monotonic across all frequencies and is best described by a single exponential function. One reason for exponential growth could be that the spectral levels (level per cycle) of the stimulus at the lower levels are at or below the activation threshold of the low-spontaneous-rate (LSR) fibers in many individuals. In mammals, the LSR fibers predominantly innervate a yet-to-be-identified interneuron in the cochlear nucleus which in turn innervate the facial motor neuron (FMN; Kobler, Guinan, Vacher, and Norris, 1992). Axons of the FMN converge on the stapedius muscle to initiate contraction (Guinan, Joseph, and Norris, 1989; Lee, de Venecia, Guinan, and Brown, 2006). When the LSR fibers are stimulated more robustly at progressively higher levels, a rapid increase in the reflex magnitude would be expected. The exponential growth could be related to the steeper slopes of the LSR rate-level function prior to saturation at very high input levels. The differences in rate-level function of the LSR fibers may also have some bearing on the differences in MEM across frequencies (Cooper and Yates, 1994; Liberman, 1978).

A frequency effect was not observed for detections. That is, all frequencies had similar intercepts and slopes for detection. Although at the outset it may seem inconsistent with the significant pairwise difference for almost all frequencies for MEM, detection rate and magnitude of change need not be highly correlated. So long as there is a significant change in the reflectance, i.e., the change is above the reflex threshold with sufficient SNR, the actual magnitude of this reflectance change should not affect detection any further. In other words, above the reflex threshold, it is the SNR that is critical for detection, not the magnitude. This result suggests that despite the small MEM at higher frequencies, the reflex nonetheless exists and engenders a small but statistically significant change in reflectance. It should, however, be emphasized that statistical significance does not necessarily imply clinical significance. Despite the reflex being present, its functional and clinical consequence may vary with magnitude. These aspects may have implications for MOCR assays aiming to avoid MEMR influence.

### IV.F. Level and frequency effects on MEMR kinetics

The time series in the present paradigm did allow us to deduce onset latency and time constants from the two-term exponential fit. The onset latency (median = 80 ms) in our data is consistent with prior reports in humans when a change in the middle ear impedance was measured using an acoustic probe. Clemis and Sarno (1980) reported a mean latency of was 105 ms (SD = 18 ms) for a 1 kHz tonal elicitor. Dallos (1964) reported latencies between 40 and 160 ms for narrowband elicitors monitored using probe tones from 250 to 1500 Hz. Qiu and Stucker (1998) reported mean latencies for various elicitors ranging from 118 ms (SD = 30 ms) for low frequency band noise to the longest at 157 ms (SD = 25 ms) for clicks presented at 50 Hz. However, Neergaard and Rasmussen (1966) reported a median latency of 17 ms when the MEMR magnitude was measured using electromyography.

Prior studies report the latency of MEMR but not time constants. Although Dallos (1964) and Hung and Dallos (1972) reported reflex kinetics similar to those observed in our study, they did not report any time constants. Therefore, it is not possible to draw direct comparisons between our data and prior studies. A frequency effect for *tau_f_* was unexpected because, assuming the change in reflectance arises from the stapedius muscle contracting, it would be expected to evolve synchronously across frequencies over time. Difference in the reflex magnitude across frequency is an unlikely candidate because as demonstrated in Section III.C, the slopes of magnitude change is not statistically significant across frequencies. In addition, we reanalyzed our time series data (not shown) by normalizing the time series Δ to be between 0 and 1 for all frequencies and levels. This transformation did not change our results. However, this result may in part be reconciled with the differences either in the shape and/or the magnitude of the time series data as discussed in Section IV.C. Such frequency differences in the time course across frequencies may be conjectured to arise from the frequency dependance of complicated motion of the tympanic membrane (Cheng, Hamade, Merchant, et al., 2013).

Although both time constants demonstrated a level X frequency interaction, they were in opposing directions. Increasing level increased *tau_f_* for low frequencies relative to high frequencies, however, increasing level decreased *tau_s_* for low frequencies. It should be emphasized that these interactions were largely driven by the difference in time constants between 1 kHz and 3.2 kHz for *tau_f_* and between 1 and all other frequencies for *tau_s_*. These interactions, although perplexing, can be explained by the marked differences in the shapes of the time course between low and high frequencies. As seen in Fig. 2, low frequency time course curves have a prolonged ‘rise time’ compared to their high frequency counterparts, this difference increases as the level is increased.

This difference may explain the *tau_f_* level X frequency interaction. Following this prolonged onset, however, the low frequencies appear to reach a steady state, likely resulting in shorter *tau_s_* times. The high frequencies, however, although ‘rise’ faster, their non-monotonic behavior can be blamed for the time course not reaching a steady state. The two-term exponential models fit these data are more likely to predict a longer (re: 1 kHz) *tau_f_*. It can again be conjectured that these frequency X level effects likely arise from the complicated tympanic membrane motion and possibly the contraction and relaxation dynamics of the stapedius muscle rather than neural factors for reasons discussed earlier (Section IV.C). It is also possible that the tau estimates are noisier than the MEM estimates, leading to variable results even between two time constants derived from the same fit. Further studies with more control over the middle ear muscle action, e.g., animal models, may be able to answer these questions in detail. Whatever may the reason for level X frequency interaction of time constants, the focus of the present study, and the advantage of monitoring the time series, nevertheless, lies in its ability to suggest whether an observed change in the stimulus level is MEMR-mediated or not (explained further in ‘Implications for MOCR assays’).

### IV.G. Implications for MOCR assays

One of the main goals of this study was to introduce a time series-based method of MEMR detection, which could also be used to minimize MEMR influence on MOCR assays. Our findings suggest that the MEMR is likely to be activated when using click trains which also activate the MOCR. This finding is consistent with previous studies for both the MEMR (Feeney and Keefe, 1999; 2001) and the MOCR (Guinan et al., 2003; Zhao and Dhar, 2012; Abdala et al., 2010; Goodman et al., 2013; Boothalingam and Purcell, 2015; Boothalingam et al., 2018) and as such, provides confidence in our methods. Perhaps the major contribution of the present study comes from its use of a time series. Although there are several methods already available that can detect the presence of a change in the stimulus level between two conditions (e.g., resampling techniques), uncertainty as to whether the observed stimulus change is truly due to MEMR still persistent in these methods. This is because, while a statistically significant change in stimulus level can occur due to the MEMR, it can also occur due to probe slippage, change in middle ear pressure during the course of the experiment, and other physiological/non-physiological factors. Conventional methods that reduce data to only two conditions (with and without noise activator) cannot readily differentiate MEMR-from non-MEMR-mediated change in the stimulus. However, we speculate that the combination of significant and the characteristic exponential change in stimulus level may be useful in more certainly attributing the change to the MEMR. Knowing whether the observed stimulus change is due to MEMR can be critical to the decision making on MOCR measurements. For example, if a change is due to the MEMR, the stimulus level may need to be lowered. In contrast, if the change is due to other factors, these may need to be addressed followed by a retest.

Monitoring the stimulus to detect MEMR activation is complicated by the reactive component of impedance varying as a function of frequency. Our findings corroborate power reflectance data of Feeney and Keefe (1999; 2001), in that, at lower frequencies (between 0.9 and 2 kHz), the impedance changes due to MEMR activation causes a reduction in stimulus level in the ear canal. In contrast, for frequencies between 2 and 3.2 kHz, the MEMR causes a relative increase in stimulus level in the ear canal. This frequency specific effect means that simple averaging across frequency of the stimulus waveform may underestimate the change in stimulus level due to MEMR activation as opposing direction of changes across frequencies may partially cancel out. Therefore, it would be useful to separate the stimulus into multiple frequency bands to examine MEMR effects.

In a simple comparison in our study (Fig. 8), ∼94% of our participants had higher MEMR thresholds when estimated using a clinical tympanometer. This discrepancy is despite the fact that broadband noise was used to elicit the MEMR in the clinical tympanometer, a superior MEMR elicitor relative to clicks (Guinan et al., 2003; Popelka, Karlovich, and Wiley, 1974). Alternatively, it is possible that our bilateral click presentation was more potent in activating the MEMR relative to the contralateral broadband noise. These differences cannot be reconciled when a different instrument and/or MEMR elicitor is used to estimate the MEMR and the MOCR. Therefore, for OAE-based assays of the MOCR, it is critical that the presence of the MEMR is *not* based on MEMR thresholds obtained from clinical tympanometers. We emphasize that monitoring the OAE-evoking stimulus is currently the optimal solution to sensitive monitoring of MEMR activation, in keeping with previous recommendations (Guinan, 2010; Zhao and Dhar, 2012; Mertes and Goodman, 2015; Boothalingam and Purcell, 2015).

Given that the MEMR appeared to be activated even at fairly low levels in some individuals, a natural question is whether it is possible to estimate MOCR effects without MEMR influence? A very small but statistically significant systematic change in the time series, presumably caused by the MEMR, does not necessarily have a measurable influence on MOCR measurements. So, this remains a pertinent question, especially for interpreting MOCR shifts. Further, the low thresholds reported in this study may not apply to other stimuli (such as broadband noise MOCR activators or MOCR activators presented contralaterally). To settle this issue fully, the magnitude of the MEMR at which the MOCR measurements demonstrate significant influence of the MEMR must be ascertained in further studies, possibly using animal models. This is a non-trivial problem because both the MOCR and the MEMR magnitude grow similarly with stimulus level and have similar time courses. Until such data are available, it is best to minimize the probability of activating the MEMR in MOCR assays.

Along with a reliable MEMR detection method, already available methods can be used to minimize the probability of evoking it. Very slow click rates (e.g. 5 Hz), low click levels, and a low contralateral noise elicitor level can be used. In addition, estimating the MOCR at multiple elicitor levels will allow room for ignoring levels with MEMR contamination. However, when low click rates/levels or low elicitor levels cannot be used the frequency-specific stimulus changes potentially provide a silver lining for MOCR assays. Our data, along with the data of Feeney and Keefe (1999; 2001), shows that the ear canal and the middle ear impedance has a magnitude zero-crossing between 1.6 and 2 kHz [see Fig. 5 panels MEM-IO and MEM-Tx]. This zero-crossing, or the transition from a reduction in stimulus level to an increment, appears to be consistent in all 20 participants in our study. It is possible that measuring MOCR magnitude changes in the 1.6 and 2 kHz region could minimize contamination by the MEMR. An individualized approach to identifying the zero-crossing frequency and capitalizing MOCR measurements close to this frequency has been reported (Goodman et al., 2018). This approach, however, assumes that the presence of MEMR at frequencies outside of the 1.6 to 2 kHz region has minimal influence on the MOCR estimated between 1.6 and 2 kHz. This assumption may not be valid. Although MOC neurons themselves are frequency specific, their dendrites within the cochlea branch extensively (Brown, 2014). Given this branching of the MOC fibers, the MOC inhibition of OAEs between 1.6 and 2 kHz is likely to be affected by inputs to the MOC neurons from other frequency regions. Testing this approach empirically in a future study with two groups of individuals, with and without MEMR, will provide evidence for MEMR influence on the MOCR in this zero-crossing frequency region.

Another possible avenue to minimize MEMR influence is to exploit the faster adaptation of the MEMR at higher frequencies by focusing on MOCR measurements at higher probe frequencies. Again, the same caveats that apply to frequency effects above also apply here. In addition, considering our time course only extended up to ∼1 s, it cannot be ascertained whether the adaptation was complete, if it only reached a threshold and continued to exist, or if it crossed the zero-magnitude line and continued to increase in the other direction. Further studies with longer click trains are required to clarify this. It is evident that when the MEMR is detected in an MOCR assay it cannot be teased apart, at least using current methods. Prevention is better than cure applies to MEMR activation in MOCR assays.

## V. Conclusion

We used bilateral click trains to generate time series data, such that MEMR-mediated changes in the impedance characteristics of the middle ear could be tracked both in frequency and time. Our approach allows for (1) the possibility of bilateral activation, (2) presentation of the probe without a separate elicitor, (3) wideband stimulus reflectance monitoring, and (4) verification if that the stimulus change is caused by MEMR by tracking the stimulus-change time series. Our data, like several previous studies, suggest that the MEMR is activated at levels lower than that reported in clinical tympanometers. Based on current evidence, monitoring the OAE-evoking stimulus in a frequency specific manner for MEMR activation is the optimal way to detect the possible influence of the MEMR in MOCR assays. The frequency specific changes observed in middle ear impedance may suggest avenues for circumventing or minimizing the effects of the MEMR on MOCR assays but need further investigation.

## Acknowledgements

The authors thank Kristina Broyles for collecting data and Dr. Viji Easwar for help with statistics. This study was supported by funds from the Office of the Vice Chancellor for Research and Graduate Education, University of Wisconsin-Madison, to SB.

